# Simulation of gap junction formation reveals critical role of cysteines in connexon coupling

**DOI:** 10.1101/2023.07.19.549697

**Authors:** László Héja, Ágnes Simon, Julianna Kardos

## Abstract

Despite the fundamental functions of connexin gap junctions (CxGJs), understanding of molecular mechanisms, governing intercellular CxGJ formation by coupling of connexon hemichannels (CxHCs, connexons) is still in infancy. In silico simulation of intercellular connexon coupling of two Cx43HC models, embedded in membrane bilayers (Cx43HC-HC) successfully modelled the emergence of trans-gap junctional (trans-GJ) stabilization centers (SCs). Investigating the molecular determinants shaping the HC-HC interface, we revealed that the exceptionally high number of cysteine residues located at the interface play a pivotal role in the stabilization of HC and GJ structures. Opening of the disulphide bonds between these cysteines resulted in disappearance of trans-GJ SCs in the Cx43GJ model. In contrast, the Cx43HC form was found to be consistent with open disulphide bonds. Finally, we have shown that the presence of an adjoining HC contributes to disulphide formation and consequently to the emergence of trans-GJ H-bonds.

Our results suggest that several connexon channels in vertebrates may undertake intercellular connexon coupling similarly and may bring forward to the targeting of CxGJ-specific coupling.

## Introduction

Connexin (Cx) hemichannel hexamers (HCs), also known as connexons, are embedded in cellular membranes and couple to form intercellular gap junction (GJ) channels. These channels serve as a conduit for molecular transfer between adjoining cells and also play a role in specific linkage ^1–4^. Despite the expanding knowledge on the cellular regulation, structure, operation and functions of Cx GJ channels ^5–30^, the mechanism of intercellular connexon coupling remains unresolved ^31^, leaving only a few initial clues. The CxHCs contain 36 Cys residues near the HC-HC interface. Primary studies revealed the principal role of disulphide bonds between these conserved Cys residues in shaping CxGJ channels by stabilizing the structure of extracellular loop motifs C(1)XXXXXXC(2)XXXC(3) and C(1)XXXXC(2)XXXXXC(3) ^32–37^. Atomic scale understanding of the role of these Cys residues, as well as other structural features in the formation of GJ channels, however, can be addressed only by *in silico* tools at present. The recent progress in the number and resolution of connexon structures (SI Table S1) allowed us to evaluate the molecular determinants of intercellular connexon coupling.

GJ channels have received attention due to their pivotal roles in various physiological and pathophysiological processes ^38–51^, such as epilepsy ^52^, Alzheimer disease ^53^ or cancer ^54–56^. However, the lack of Cx subtype-specific inhibitors for these channels has hindered their exploitation in pharmacological strategies. Since GJ channels have no specific binding sites or endogenous antagonists, the general approach to design specific inhibitors was to mimic Cx sequences of the EL1 and EL2 extracellular loops, which are thought to be involved in HC-HC coupling. Mimetic peptide inhibitors, however, have never demonstrated to be Cx-specific. Using molecular docking, we also showed that they do not bind to the sequence where they were designed to ^57^. Instead of mimicking consecutive sequences of extracellular loops, we consider that Cx-and Cx-subtype specificity can be achieved by identifying the residues playing primary role in trans-gap junctional (trans-GJ) interactions, paying special attention to protein stabilization centers (SCs), hubs of interactions that build up the HC-HC interface ^58^.

In the current study, we aimed to investigate the structural determinants of GJ formation, including the role of Cys residues near the HC-HC interface. We used the homomeric Cx43 channel as a connexon prototype. Cx43GJ channels are abundant on astrocytes and play a crucial role in the development and regulation of neural circuit function and animal behaviour ^52,53,59–64^. We created a Cx43HC model based on the experimental connexon structure Cx31.3HC (PDB code: 6l3t; SI Table S1, SI Fig. S1A)^65^. To simulate HC-HC coupling, two membrane bilayer-embedded Cx43HCs were paired according to the architecture of the homomeric dodecamer Cx26GJ channel (SI Fig. S1B) and the HC-HC coupling process was simulated by placing the two HCs at varying distances (Cx43HC-HC).

## Results

### Simulation of Cx43 connexon coupling: the development of trans-GJ SCs

To understand GJ formation from HCs and to identify regions and structural motifs playing pivotal roles in the process, we first modelled Cx43HC based on the high-resolution cryo-EM structure of Cx31.3HC (SI Tables S1), the only published experimental solo HC structure ^65^. It is to mention that, although an experimental Cx43 HC structure was published recently ^66^, it was obtained from GJ proteins, therefore it rather represents the HC as it exists in the full GJ form. Recently, a solo Cx43 HC structure was also deposited in the protein database ^67^. To compare our model to these structures, we determined the residue-level RMSD values between the model and the experimental structures. The model showed comparable or even higher similarity to the experimental structures than the two experimental structures to each other (SI Fig. S2), especially in the HC-HC interface region (residues 55-58 and 194-196). In addition, we created a Cx43GJ homology model using the experimental Cx26GJ structure as a template ^57,58,68^. The generated Cx43GJ model also showed high similarity to the recently published Cx43 GJ structures ^66,67^ (SI Fig. S2). To simulate the coupling of HCs to form GJ, we embedded two Cx43HC models into explicit membranes and positioned them face-to-face according to the arrangement observed in the Cx43GJ model (Fig. 1A).

**Figure 1.**
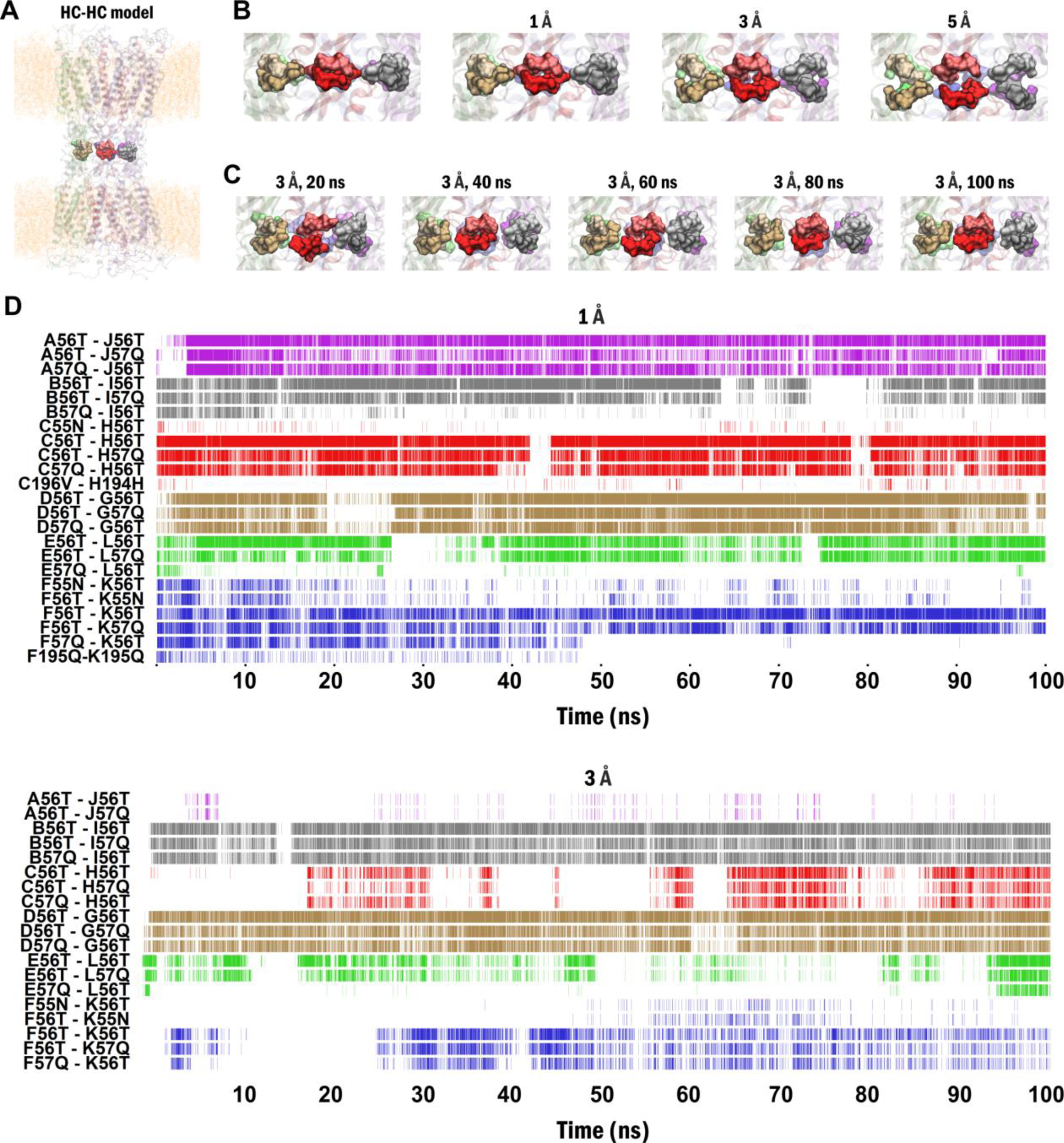
Spatio-temporal coordination dynamics of Cx43HC-HC reveal the build-up of trans-GJ SCs as an indicator for HC-HC coupling. (A) Large-scale side view of two membrane embedded Cx43HCs, modelled based on the cryo-EM Cx31.3HC template (PDB code: 6l3t; SI Table S1) ^65^, were joined up in the architecture of the homomeric dodecamer Cx26GJ channel (PDB code: 2zw3; SI Table S1) ^68^ (Cx43 HC-HC). Residues 55N-56T-57Q of pre-and post-GJ subunits A-J, B-I, C-H, D-G, E-L and F-K are highlighted light-and dark purple, grey, red, brown, green and blue, respectively. (B) Close view of the interface residues 55N-56T-57Q of the Cx43 HC-HC model show gradual disintegration of the trans-GJ interface with increasing distance between HCs. (C) Presence of specific trans-GJ SCs during 100 ns MD simulation show rebuilding of trans-GJ interactions after moving away HCs to 3 Å distance. SCs defined at the A-J, B-I, C-H, D-G, E-L and F-K interfaces are highlighted in purple, grey, red, brown, green and blue, respectively. Trans-GJ SC pairs are designated as subunit name + residue number + 1-letter residue name. (D) Close view of the HC-HC interface residues during the 100 ns MD simulation after moving away HCs to 3 Å distance demonstrate rebuilding of trans-GJ interactions. Note the development of trans-GJ coupling between subunit pairs C-H (light and dark red).

After setting up the initial paired Cx43HC (Cx43HC-HC; Fig. 1A), we moved the two HCs away to distances of 1, 3 and 5 Å to model different stages of the coupling process (Fig. 1B). All models were subjected to 100 ns MD simulations and we continuously monitored the appearance of SCs, previously shown to build up the HC-HC interface. ^58^ When HCs were positioned at their original locations or at a distance of 1 Å (*see* Methods section), the HC-HC interface formed from residues 55N-56T-57Q ^58^ were stable at the beginning of the simulation (Fig. 1B). Increasing the distance to 3 Å and 5 Å introduced a significant and increasing gap between the two HCs (Fig. 1B). During the 100 ns simulation, the HC-HC interface was observed to be reappeared when the original HC-HC distance was set to 3 Å (Fig 1C).

To quantitatively address the HC-HC coupling strength, we investigated the appearance and progressive emergence of SCs involving residues from the two adjoining HCs (trans-GJ SCs), because SCs are expected to have major contribution to the stability of the full GJ channel. Gradual build-up of trans-GJ SCs were considered as an indicator for successful coupling between the two HCs. In the initial model, in which HCs were arranged according to their original position, observed in the Cx43GJ model, several trans-GJ SCs could be identified between the internal 55N-56T-57Q-58Q (major) and external 194H-195Q-196V (minor) EL1 and EL2 residues, forming the interface between the two opposing HCs (Fig. 2A). When the two HCs were positioned at 1 Å distance, these trans-GJ SCs remained intact (Fig. 1D). However, once the distance was increased to 3 Å, many of these trans-GJ SCs disappeared, suggesting a weakened (i.e. not-yet fully developed) coupling. Importantly, several trans-GJ SCs was found to be re-established during the 100 ns simulation (Fig. 1D), demonstrating that the MD of pre-positioned single HCs can be an appropriate model of HC-HC coupling and GJ formation.

**Figure 2.**
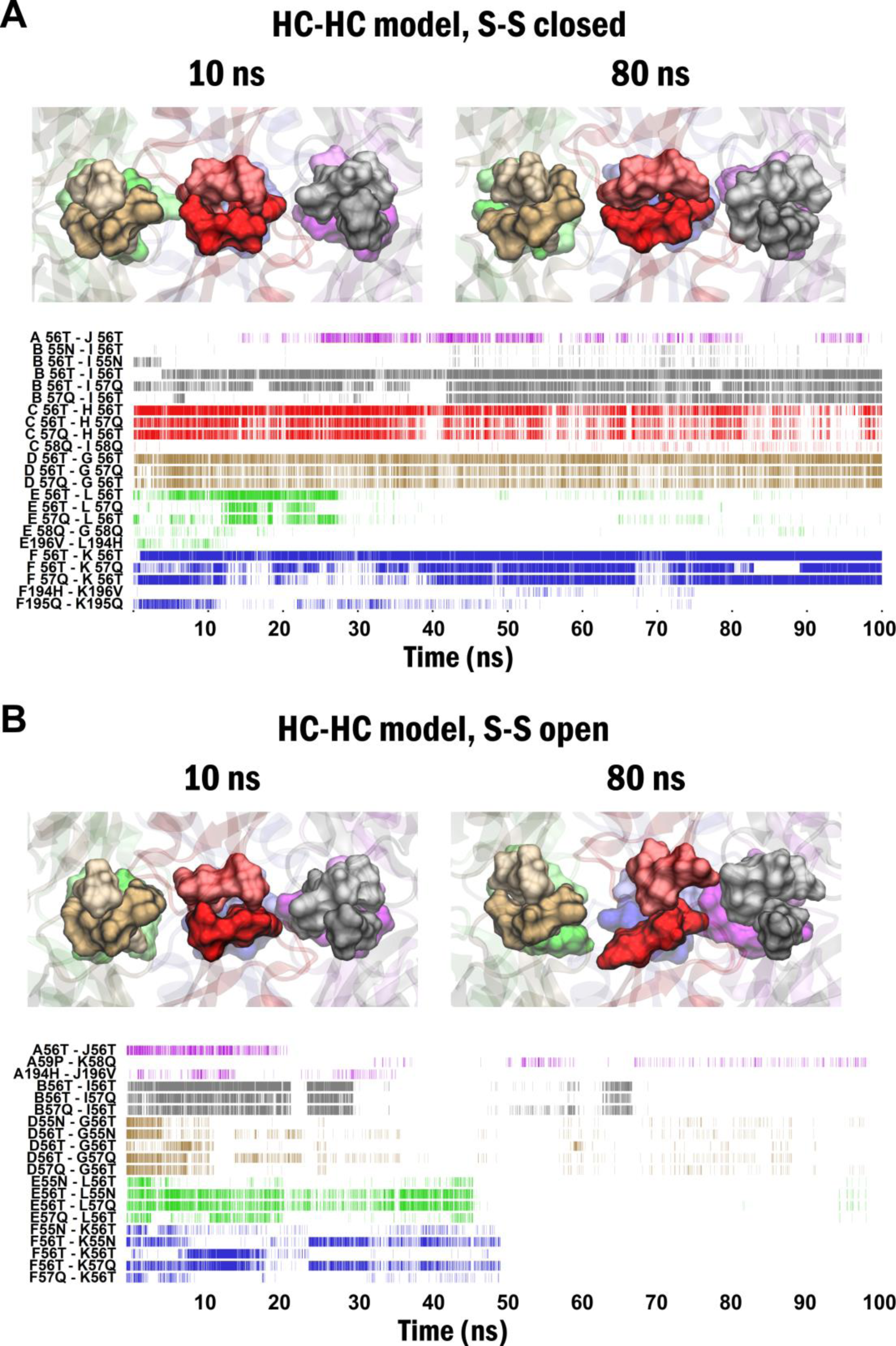
Distinguishable spatio-temporal coordination dynamics of Cx43HC-HC under closed *versus* open disulphide bonds preconditions. (**A**) Close view of the interface residues 55N-56T-57Q-58Q (top) and presence of specific trans-GJ SCs during 100 ns MD simulation (bottom) in the Cx43 HC-HC model in the closed disulphide configurations. (**B**) Close view of the interface residues 55N-56T-57Q-58Q (top) and disappearance of specific trans-GJ SCs during 100 ns MD simulation (bottom) in the Cx43 HC-HC model in the open disulphide configurations. Residues of trans-GJ SCs 55N-56T-57Q in pre-and post-GJ subunits in A-J, B-I, C-H, D-G, E-L and F-K pairs are highlighted purple, grey, red, brown, green and blue, respectively. Trans-GJ SC pairs are designated as subunit name + residue number + 1-letter residue code.

We were able to capture the appearance of trans-GJ SCs at both inner (channel-facing) and outer (gap-facing) GJ surfaces. The major SCs at the inner surface consisted of extracellular loop EL1 residues 55N-56T-57Q, while the minor SCs at the outer surface were made of the EL2 residues 194H-195Q- 196V of each paired Cx43HC-HC subunits. Interestingly, increasing the HC-HC distance to 5 Å resulted in irreversible and almost complete loss of trans-GJ SCs, leaving only.minor trans-GJ SCs between the bulky 194H and 196V residues to persist (SI Fig. S3).

By investigating what structural motifs may contribute to the proper orientation of these residues, we noticed that 55N-56T-57Q and 196V are associated with the 54C(1)-198C(3) disulphide bridge, while the sequential trans-GJ SC residues 194H-195Q conjoin the 61C(2)-192C(2) disulphide bridge (SI Fig. S4A). Some of these SCs correspond to H-bonding (51A “O”-201S “OG”; 53R “O”–199F “N”; 55N “N”-197D “O”; 58Q “NE2”-193P “O”) while others represent Van der Waals contacts between carbon atoms. Three-dimensional structure of the HC-HC interface (SI Fig. S4B) explains quasi-mirror arrays of tetragonal SC patterns, apparently oriented by cystine-linked trans-GJ SCs and H-bonding interactions (red lines) (SI Fig. S4C). Beside trans-GJ contacts, the pattern of cystine-linked intra-HC SCs is shown by SI Fig. S4D. The unique 65C(3) centred three dimensional SC pattern exhibits GJ contacts within and between EL1 and EL2 loops.

### Opening of disulphide bonds abolishes trans-GJ SCs

Given the abundance of disulphide bonds in the neighbourhood of the trans-GJ SCs and the ability of cysteine thiols to spontaneously transform to cystine disulphides ^69–73^, we investigated whether dynamic S-S bond opening and closing may contribute to the transition from HC to GJ structures. To assess this hypothesis, we opened disulphide bonds and applied 100 ns MD dynamics on Cx43HC-HC model at zero or intermediate (3 Å) distance between the two HCs. The assessment of spatio-temporal dynamics demonstrated that the opening of disulphide bonds resulted in the disappearance of all trans-GJ SCs during the 100 ns MD simulation when HCs were positioned at zero distance (Fig. 2). At the intermediate distance (3 Å), no trans-GJ SCs were developed during the 100 ns MD, in contrast to the closed disulphide condition (Fig. 1C). Notably, inter-subunit SCs, stabilizing connexin-connexin coupling within HCs remained unchanged (SI Fig. S5), demonstrating that opening of disulphide bonds specifically affected the HC-HC interface interactions. In summary, these results show that opening of disulphides in the HC structure prevents GJ formation and may keep connexins in the HC form.

### Are disulphide bonds open in the HC structure?

The presence of disulphide bonds can only be assumed from the experimental structures. Although all currently available connexin structures contain disulphide bonds, this could be a result of experimental conditions used in structure determination. We have shown that trans-GJ interactions abolish after opening of disulphide bonds, in accordance with experimental data ^32,74^. However, HCs can be fully functional even when all extracellular Cys residues were transformed to thiols ^75^, suggesting that disulphides are crucial only in the GJ, but not in the HC form ^74^. To explore whether disulphides are closed or open in the solo HC, we compared the Cx43HC model based on the Cx31.3 template that represent the solo HC with the A-F subunits of the Cx43GJ model that represent the structure of a HC as it exists in a full GJ. Finally, we also compared the Cx43HC model to the Cx43HC-HC model to assess the effect of constraints induced by the close presence of the opposing HC. All structures were subjected to 100 ns all-atom MD simulations with disulphide bonds kept closed or opened at the beginning of the MD run.

We found that in the Cx43GJ model, significantly less conformational changes are required to reach a steady-state structure in the closed disulphide configuration compared to the open disulphide configuration, suggesting that closed disulphide bonds are, indeed, represent the typical state of Cx43 GJ (Fig. 3A). In contrast, in the Cx43 solo HC models, despite of the higher degree of freedom, the open disulphide configuration approaches the steady-state structure more rapidly, implying that open disulphides may be the physiological conformation of the solo HC (Fig. 3B). However, presence of the opposing HC in the Cx43HC-HC model results again in significantly less conformational change in closed disulphide configuration compared to the open one (Fig. 3C). Comparisons of top views of open disulphide Cx43HC-HC, solo Cx43HC and Cx43GJ models indicate that the open disulphide Cx43HC-HC is markedly different from the solo Cx43HC, instead it is similar to Cx43GJ (Fig. 3D). These data suggest that open disulphides may be the more appropriate condition under which solo HCs exist, but presence of the opposing HC may shift the structure to the Cx43GJ architecture, a state in which closed disulphides are more favored.

**Figure 3.**
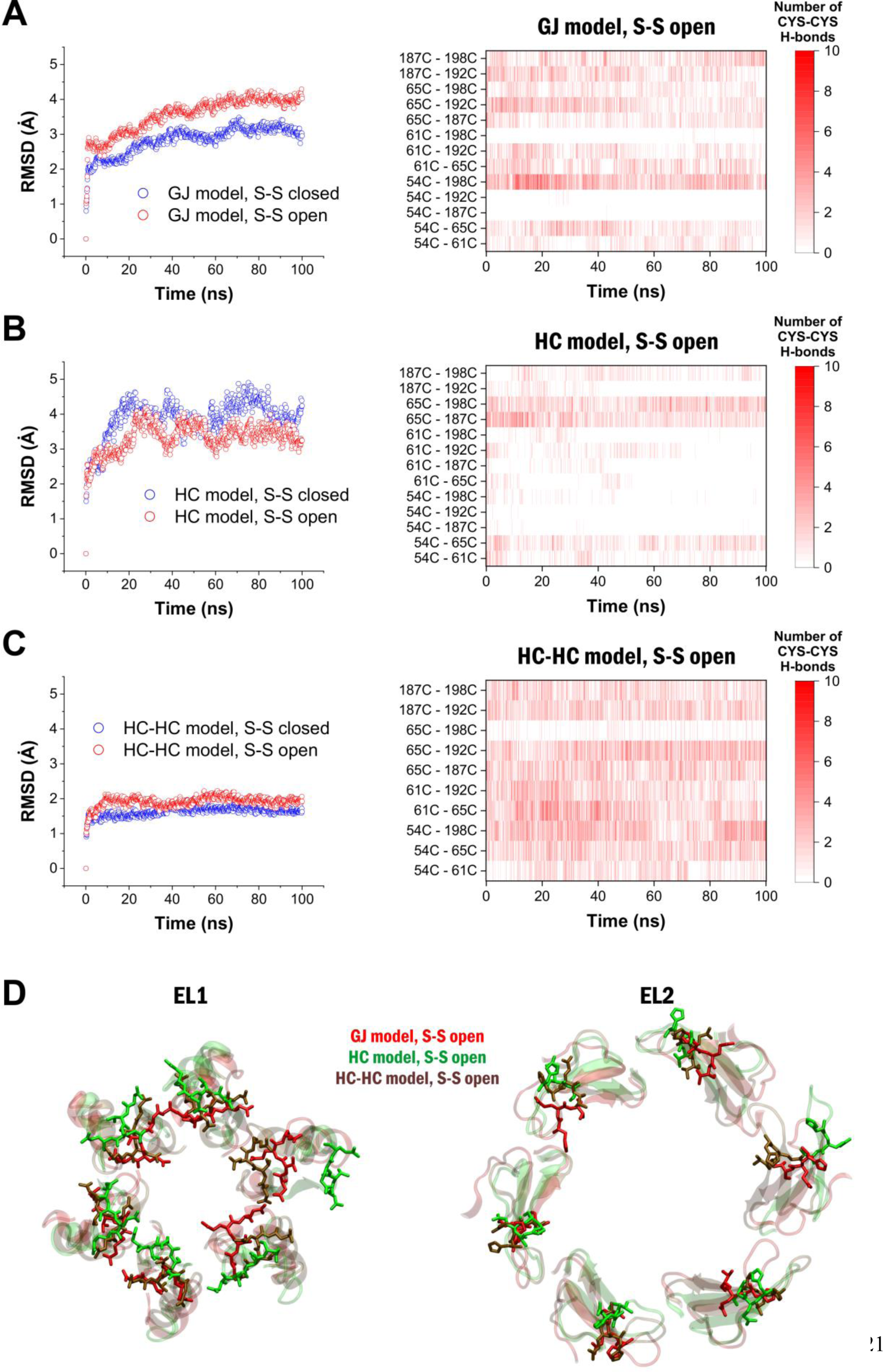
Solo HC structure may be better represented by open disulphide bonds. **(A)** RMSD changes of the extracellular region compared to the initial structure (left) and number of specific H-bonds between Cys residues during the 100 ns MD (right) in the Cx43GJ model. **(B)** RMSD changes of the extracellular region compared to the initial structure (left) and number of specific H-bonds between Cys residues during the 100 ns MD (right) in the Cx43HC model. **(C)** RMSD changes of the extracellular region compared to the initial structure (left) and number of specific H-bonds between Cys residues during the 100 ns MD (right) in the Cx43HC-HC model. In order to compare HC model with GJ and HC-HC models, only the number of H-bonds on the A-F subunits were counted. Theoretically, 2 H-bonds can be formed on each subunits (both Cys residues can be either proton donor or acceptor), totaling 12 H-bonds between Cys residues on all subunits. **(D)** Extracellular view of the EL1 (left) and EL2 (right) regions of the A-F subunits of the Cx43GJ (red), Cx43HC (green) and Cx43HC-HC (brown) models with open disulphide bond conditions at the end of the 100 ns MD runs. Extracellular loops are shown in cartoon representation, interface residues (55-58 and 194-196) are shown in stick.

In addition, we also analyzed what Cys-Cys H-bond interactions show up in the open disulphide configurations during the simulations to estimate the possibility of different disulphide bond formations. According to this analysis, in the Cx43GJ model, many different potential disulphide bonds can be formed throughout the MD run, including the 54C(1)-198C(3), 61C(2)-192C(2) and 65C(3)-187C(2) disulphide bonds, present in the experimental Cx26 structure (Fig. 3A). These data show that despite opening the disulphide bonds at the beginning of the MD run, they can be easily rearranged. In contrast, opening the disulphide bonds in the Cx43HC model led to structural changes that opposed disulphide re-formation by keeping Cys residues far from each other to bind (Fig. 3B). In the Cx43HC model, the original 54C(1)-198C(3), 61C(2)-192C(2) and 65C(3)-187C(3) disulphide bonds are almost completely missing. The Cx43HC-HC model was found to be similar to the GJ model, enabling all the original and many other disulphide bond formations (Fig. 3C), confirming that the presence of opposing HC leads to restructuring of solo HC and favoring of disulphide formation.

Conclusively, comparison of GJ, solo HC and HC-HC models suggest that open disulphide bonds may better represent the HC form. Therefore, it is rational to hypothesize that disulphide bonds are mainly open in the HC structure, but are closed for the most part of the full GJ channel.

### Presence of free cysteine in HC prevents H-bond interactions at the HC-HC interface

Our data showed that solo HC structure is consistent with open disulphide bonds (Fig. 3). However, opening of disulphide bonds in the Cx43HC-HC model resulted in the disappearance of trans-GJ SCs (Fig. 4B). These results are in line with previous observations showing that closed disulphide bonds are necessary for GJ formation, but are not required for solo HC function ^74,75^.

**Figure 4.**
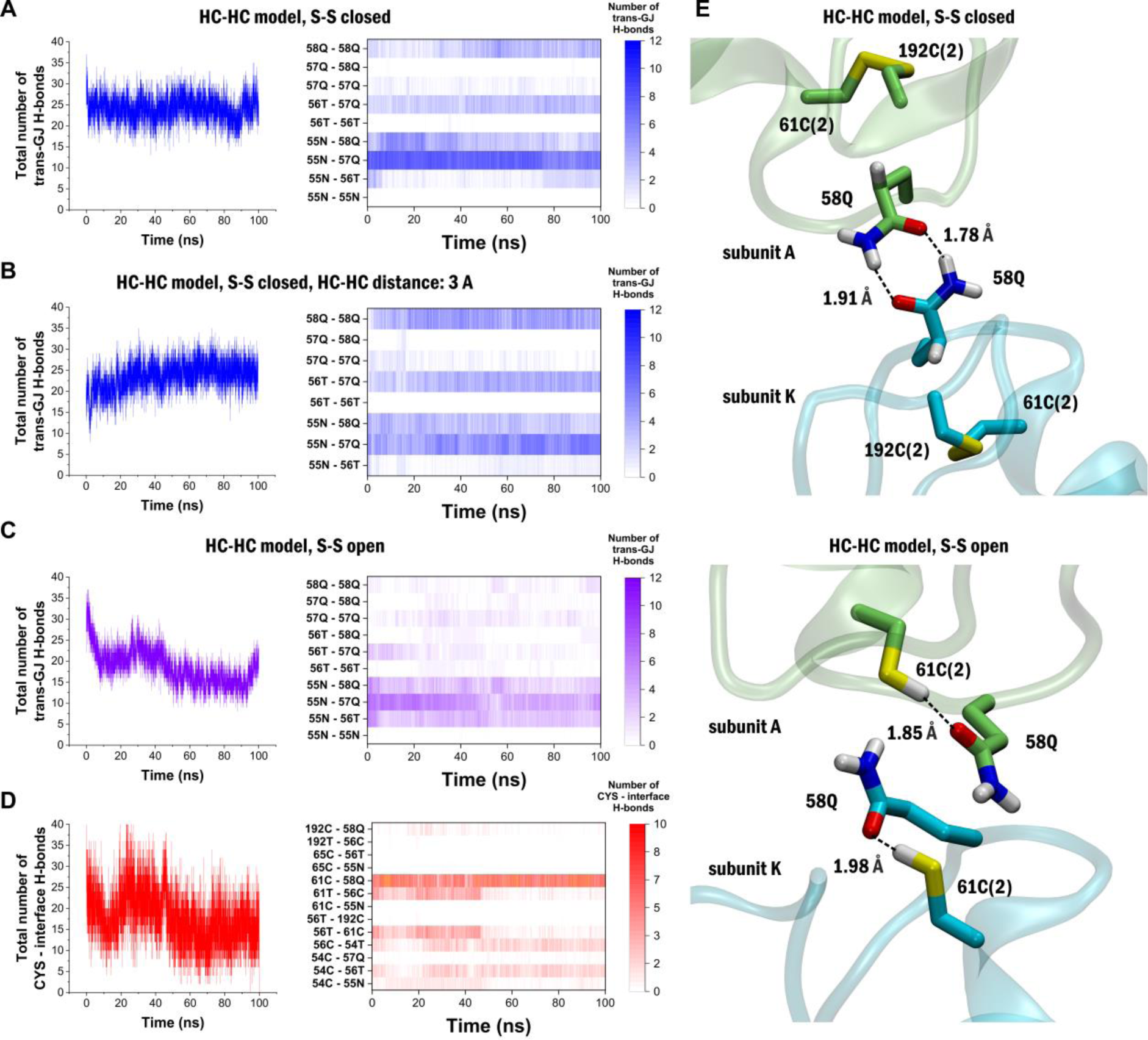
Free cysteine residues Cys54, Cys61 and Cys192 disorient residues involved in trans-GJ interactions. (**A**) Number of trans-GJ H-bonds (left) and specific residue pairs forming these trans-GJ H-bonds (right) during the 100 ns MD in the HC-HC model with closed disulphides. (**B**) Number of trans-GJ H-bonds (left) and specific residue pairs forming these trans-GJ H-bonds (right) during the 100 ns MD in the HC-HC model with HCs positioned 3 A away. (**C**) Number of trans-GJ H-bonds (left) and specific residue pairs forming these trans-GJ H-bonds (right) during the 100 ns MD in the HC-HC model with open disulphides. (**D**) Number of H-bonds involving Cys residues (left) and specific residue pairs forming these H-bonds (right) during the 100 ns MD in the HC-HC model with open disulphides. Colour intensity in A-D corresponds to the number of subunit pairs on which the specific H-bond is present. (**E**) 3D representation of the HC-HC interface shows disorientation of 58Q from the position required to make trans-GJ H-bonding. Trans-GJ H-bonds between opposing Gln58 residues are formed in the HC-HC model with closed disulphides (top). In the open disulphide model (bottom) 58Q oxygen atoms are involved in H-bonds with the free Cys residues instead of forming trans-GJ interactions.

To assess what structural changes may lie behind the transition process, we explored which residues are involved in the interactions between the two HCs. We observed that trans-GJ H-bonds are formed between residues 55N, 56T, 57Q and 58Q in the closed disulphide HC-HC model (Fig. 4A). The same interactions emerged during the MD simulation when the two HCs were initially positioned at 3 Å distance (Fig. 4B). Some of these interactions, especially those forming between 58Q residues on both HCs disappeared when the disulphide bonds were open (Fig. 4C). Analyzing the structural changes corresponding to the in silico reduction of CYS residues, we found that the free Cys residues in the open disulphide model can form intra-subunit H-bonds with various residues involved in the trans-GJ interaction, thereby weakening the HC-HC coupling potential (Fig. 4D). For example, intra-subunit H-bonding between 61C(2) and the oxygen atom of 58Q prevents the formation of H-bonds between the opposing 58Q residues (Fig. 4E). Therefore, keeping the HCs in open disulphide configuration inhibits HC-HC coupling and GJ formation.

## Discussion

Although several lines of structural information from cryo-EM and single particle analyses of Cx43 GJ channel have been reported recently ^66,67^ (SI Table 1), structural clues concerning intercellular coupling of Cx43 HCs have not been predicted until now. In this study, we simulated intercellular connexon coupling by pairing two membrane-embedded Cx43 HCs. The simulations revealed the consecutive build-up of trans-GJ SCs. By investigating the structural prerequisites of HC-HC coupling we revealed that residues of the internal 55N-56T-57Q-58Q and external 194H-195Q-196V HC-HC interface in Cx43 are oriented to the appropriate conformation by the large number of neighbouring Cys residues. By exploring the structures of Cx43HC and Cx43GJ or Cx43HC-HC models, we found that the HC form may be coherent with open disulphide bonds. These observations provide mechanistic clues that relate gap structure and coupling through the cystine-linked development of subtype-specific trans-GJ interactions.

The emergence of cysteines in SC contacts with Arg/Lys in the gap as characterized by minimum distances 3.34 Å and 3.23 Å in 65C(3)-189Arg and 187C(1)-68Lys, respectively (SI Table S2), raises the issue of connexon coupling via oxidative disulphide bond exchange ^69,70,72,73^. We suggest the cysteine thiol to cystine disulphide redox exchange of 65C(3) and 187C(1) enabled by flanking positively charged Arg or Lys ^76–80^ as being part of the cause of GJ design. It also explains the Cx redox sensor function ^15,74,81–84^.

Now the matter arises concerning assets assigning cysteine thiols to switch over cystine disulphides? In line with previous suggestion, the low pKa cysteine thiol may spontaneously oxidize to cystine disulphide at neutral *p*H generating two protons in addition to two electrons ^77^. The process involves the formation of reactive thiolate anions that oxidize to sulphonic acid thus elevating reactivity towards nearby thiol groups ^85^. Having a correlation time in the 10^-7^ to 10^-9^ s range ^86^, the rapid proton transfer to flanking Arg/Lys can further cystine disulphide bond formation. We put forward the proton-acceptor feature of Arg/Lys that enhances the redox-sensitivity of Cys along with the rate of thiol/disulphide transformation. Running MD simulations in the 100 ns time range we noticed fast fluctuations of nearest 68Lys/189Arg residue distances from Cys in the range mainly similar with 65C(3)–187C(1). The time to distance variations mostly compare with 1-5 ns disulphide exchange events simulated for 35 ns in the β3 integrin subunit ^87^.

To investigate the structural patterns and potential role of extracellular SCs in GJ formation, we visualized the connectivity of SCs in the HC-HC model, distanced at 3 Å using a graph representation of all data of all subunits during the entire 100 ns MD run (Fig. 5A). The connectivity pattern revealed that two cysteine-centred modules can be distinguished that are connected by the central node 65C(3). The two modules, stabilized by the 54C(1)-198C(3) and the 61C(2)-192C(2) disulphides, both consist of SCs involving the EL1 and EL2 loops. These modules, as well as the group of transmembrane exits of TM2, TM3, TM4 helices are connected by the 65C(3)-associated pattern of SCs (Fig. 65A). Another independently formed SC pattern comprises of the 43S-47D, 44A-47D, 44A-45W residues that support Ca^2+^ binding by 43S, 46E and 48E. A further independent SC pattern is characterized by 66Y edging 49Q and 202R, which assists positioning of TM1 and TM4 in concert with molecular changes occurring in the TM2-connected extracellular helix. Apparently, the graph derived from SC dynamics describes that two cysteine-centred groups of SCs, connected by the conserved sequence 65C(3)-66Y-67D of the extracellular helix form the “gap syntone” that guides proper lining of 55N- 56T-57Q for HC-HC coupling (Fig. 65B). The central role for 65C(3) ↔ 189R may allow the prevention of GJ design by binding guanine to 189R ^88^. Our results also suggest that GJ SC patterns emerging from coupling at the TM helix-outer membrane interfaces indicate dynamic changes in the local lipid environment. Indeed, all-atom MD simulations of cryo-EM data from the native Cx46/50GJ in lipid discs reveal the lipid-induced stabilization of the GJ channel and vice versa ^89^.

**Figure 5.**
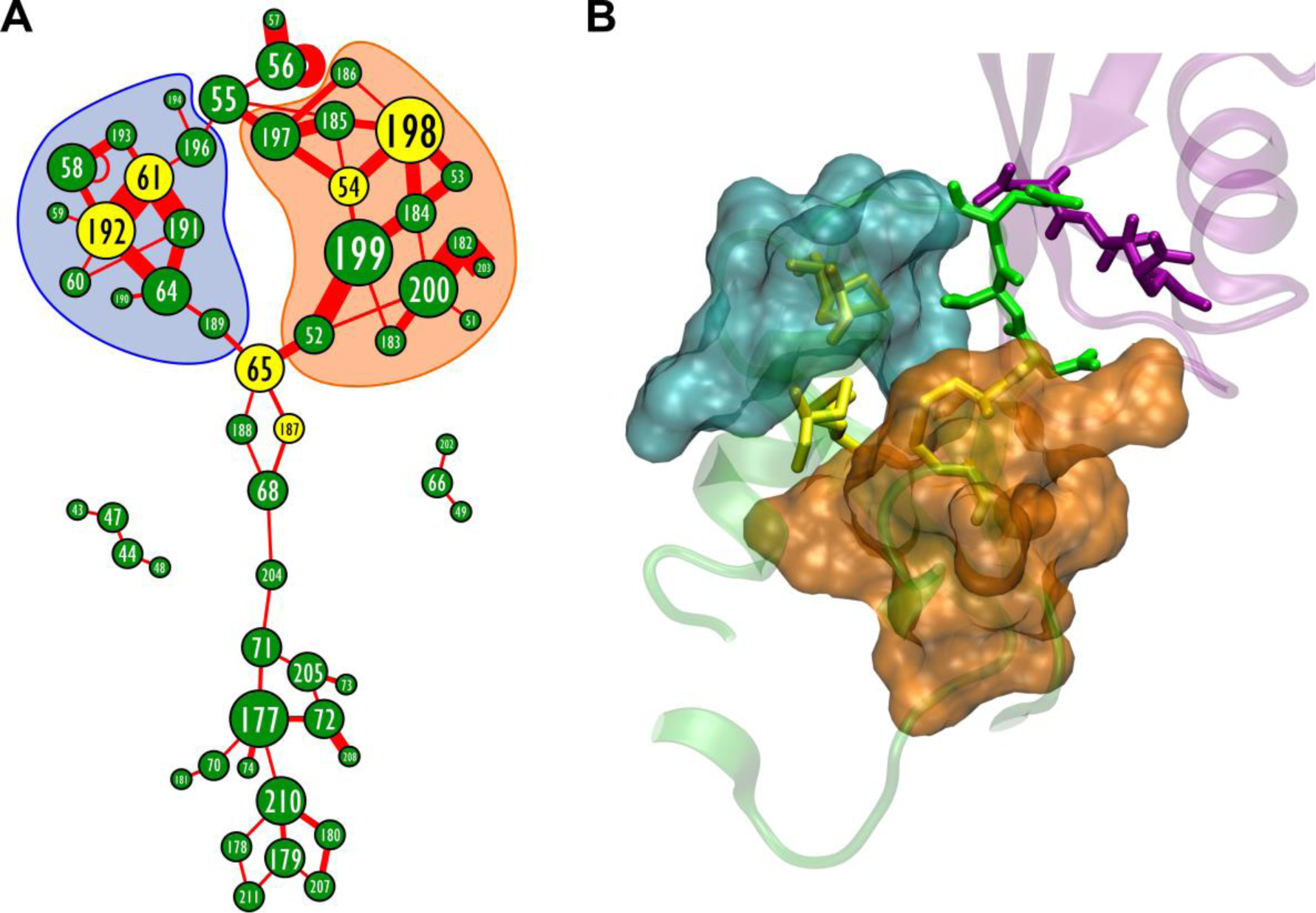
Cysteine-controlled formation of associated stabilization centers (SCs) orient Cx43HC-HC interfaces for coupling. (**A**) Graph representation of extracellular SCs in the GJ built from two opposing HCs after distance between the HCs set to 3 Å. Nodes represent residues, edges represent SCs formed between given residues. Data of all subunits during the whole 100 ns MD run is summarized into a single graph. Size of the nodes corresponds to the number of SCs the given residue is participated in. Width of the edges corresponds to the stability of the given SC during the 100 ns MD run. Cysteine residues are highlighted by yellow nodes. (**B**) 3D structure of subunits A (green) and J (purple) in the GJ built from two opposing HCs after distance between the HCs set to 3 Å. Interface residues (55N-56T-57Q) are shown in green and purple stick representations on the A and J subunits, respectively. Cysteine residues are shown in yellow stick representation. SC associations identified in **A** are shown in cyan and brown surf representations.

Our models predict that the six conserved Cys residues of EL1 and EL2 extracellular loops per Cx43 monomers directly regulate functional coupling of Cx43HCs. Now we understand the intercellular connexon coupling as redox-sensitive cysteine guided supra-molecular self-assembling ^90,91^ of interconnected SC patterns. Self-assembling can be genuinely recognized by multitude of disulphide context-specific proton donor-acceptor interactions ^39^ and associated polar and Van der Waals contact relations, where one residue principally contacts two to four residues, in addition to the attendance of Ca^2+^ complex formation ^9,25^ and likely electron shuffling ^58^.

Spatio-temporal configuration dynamics of paired Cx43HC-HC model enable the observation of atomic scale development of extracellular cystine-linked trans-GJ SCs of EL1 C(1)vvzvvv(2)zzzC(3) and EL2 C(1)zzzvC(2)vzzvvC(3) sequences, in which 100 % of C(1), C(2), C(3), >80 % of X residues (v) and <80 % of X residues (z) are conserved ^58^. In view of conserved extracellular cysteines and extensively homologous protein sequences of Cx isoforms, along with their mutations, phosphorylated and ubiquitinated derivatives ^43,44,47,92,93^, it seems likely that several homo-and heteromeric CxGJ channels in vertebrates are formed according to similar patterns. The less conserved extracellular residues do introduce subtype-specificity and may also confer non-coupling nature, as observed in Cx31.3HC ^94^.

Our trans-GJ SC interface design may serve the development of subtype-specific inhibition of intercellular connexon coupling by targeting the build-up of trans-GJ SCs. It seems feasible by triggering trans-GJ SC site-specific bonding of inhibitory ‘substrate’ (S) by adding cysteine-targeted electron withdrawing group (EWG; Michael acceptor) ^95–97^ through an actual linker (L). For example, the designated EWG-L-S blueprint expects Cx43 subtype-specific inhibition by 55H-56T-57Q isosteric non-peptide S derivatives. Our perspective also anticipates that the EWG-L-S design shall overcome potential off-target interactions ^98,99^ by enhancing the intrinsic EWG reversibility ^95^. Also, the fine tuning of span and flexibility of L can serve the Cx GJ selectivity of EWG-L-S inhibitors. However, the specific inhibition of HC-HC coupling in the molecular mechanisms of actual pathologies could be the appropriate approach to elicit side-effects. For example, decoupling of astrocytes may induce the impairment of synaptic plasticity and spatial learning memory ^100^. We place the issue in the context of cell-specific drug delivery ^101^.

## Methods

### Generation and molecular dynamics of Cx43HC model

The Cx43HC was prepared using the high-resolution cryo-EM Cx31.3HC structure as a template (PDB code: 6l3t) ^65^. Swiss-Model generated an alignment between the target and the template (SI Fig. S1A) and built the 6l3t structure-based model. After a short energy minimization, we obtained the Cx43HC model. This was submitted to the “Positioning Proteins in Membrane” (PPM) server of the “Orientations of the Membranes in Proteins” (OPM) database ^102^ to predict the trans-membrane regions: 20-46, 74-94, 154-176, 205-226, and to rotate the protein parallel to the z axis. The Cx43HC model was then subjected to 100 ns MD. In detail: The model was loaded to the workspace of Maestro-Desmond (DE Shaw Research) ^103^. The protein preparation wizard was invoked and pre-processing was performed with the “create disulphide” option checked or unchecked depending on the simulation conditions. This option ensures that Cys residues remain in disulphide-bonding state all over the simulation (referred to as “S-S closed condition”), or Cys residues are present as thiols (SH) (referred to as “S-S open condition”) Also, hydrogen bonds were added during pre-processing. A predefined membrane was added and “placed on the pre-aligned structure”. Since the predefined membrane is positioned in the *x-y* plane, the OPM-rotated protein was aligned accordingly, perpendicular to the membrane. The structure was subsequently loaded into the Desmond module. Temperature and pressure were kept constant at 300K and at 1 atm pressure, respectively (NPT condition). Simulation time was set to 100 ns, the recording interval was set to 10 ps, so altogether 10 000 frames were collected.

### Generation and molecular dynamics of Cx43HC-HC model

Two copies of the solo Cx43HC model (10824 atoms each) were taken. One copy was aligned to chains A-F of the template Cx26GJ channel with closed cystine disulphide bonds (PDB code: 2zw3; SI Table S1) ^68^ with Pymol. The other copy was aligned to chains G-L of the same template. Prior to aligning, the second copy was renumbered, to start from atom No. 10825, to provide continuous atom numbering in the newly generated dimer. This resulted in a raw model of the coupled Cx43, in which several interface residues were positioned in a clashing distance of less than 2 Å. Therefore, a short minimization was performed in Maestro on all atoms, using the “minimize all” command. Subsequently, the model was submitted to OPM, and the Cx43HC-HC model was prepared in Visual Molecular Dynamics (VMD), because VMD allows the application of two membrane bi-layers per protein ^104^. First, to generate protein structure files (psf) the structure was split into 12 individual chains. Disulphides were set between 54C(1) and 198C(3), 61C(2) and 192C(2) along with 65C(3) and 187C(1) using the DISU patch of VMD. Also, an open version was prepared, without creating disulphide bonds. The 2 x 6 apposed subunits were then combined into a single file and two POPC membrane bilayers (150 x 150 Å each) were generated by the membrane builder plugin. Both HCs were embedded in membrane and the whole system was solvated as described previously ^58^. The “keep water_out” tcl script was modified in-house to be appropriate for two membranes ^58^. Simulation time was set to 100 ns, the recording interval was set to 10 ps, so altogether 10 000 frames were collected.

### Generation and molecular dynamics of Cx43HC-HC model with varying HC-HC distance

The paired Cx43HC-HC model was taken as a starting structure. The protocol was the same as above until the first POPC lipid was put on one HC (chains G-L) and a combined lipid-protein pdb file was saved along with its protein structure (psf) file. After this stage, chains A-F were selected and moved by 1 Å in the z direction using the “moveby” command of VMD that moves each of the selected atoms by the given vector offset. After lifting all atoms of Cx43HC by 1 Å, all atoms of the POPC lipid membrane around the Cx43HC were also moved by 1 Å in the z direction and saved together with the HC containing chains A-F. Then the protocol continued identically as described above. To follow the effect of HC-HC distance in a stepwise manner, three different distances were introduced: 1 Å, 3 Å and 5 Å. Disulphides were kept closed in all cases and the simulation lasted for 100 ns.

### Generation and molecular dynamics of Cx43GJ model

The Cx43GJ model was built based on the Cx26 structure ^68^ as described before ^58^. Briefly, an initial model was built by Swiss-Model, which was submitted to OPM to predict TM regions. Subsequently, the full Cx43 GJ model was placed in two POPC membrane bilayers as in the HC-HC model. The Cx43 GJ model was prepared with S-S closed and in S-S open conditions as was the HC-HC model. MD was performed as above for 100 ns.

### Determination of stability centers

In order to identify SCs, MD trajectories from NAMD or Desmond were imported into VMD and individual frames at 10 ps interval were exported as pdb files. After adding sequence residues (SEQRES) data to the pdf files, SCs were identified using the SRide server ^105^. Extracellular SCs were ered by selecting SCs containing at least one extracellular amino acid (EL1 residues 47-73 or EL2 residues 177-203) and appearing in at least 2 % of the total running time (corresponding to 1 ns).

### Calculation of RMSD values

In order to calculate RMSD values, protein models in all frames of an MD simulation were aligned to the first frame (t = 10 ps) using the membrane segment of the protein (residues 21-46, 74-93, 156-276 and 204-229) as a base for alignment. After the alignment, RMSD changes of the extracellular part (residues 47-73 and 177-203) were calculated for all frames using VMD. RMSD values between our model and the recently published Cx43 structures were also calculated for the extracellular part (residues 47-73 and 177-203) using VMD.

### Determination of H-bonds

Trans-GJ H-bonds were identified between residues 55-58 using the following donor atoms: backbone “N”, Asn “OD1”, Thr “OG1”, Gln “NE2” and the following acceptor atoms: backbone “O”, Asn “OD1”, Thr “OG1”, Gln “OE1”. Trans-GJ H-bonds were identified when the distance between donor hydrogen atoms and acceptor heavy atoms were below 2.5 Å. H-bonds between Cys residues and residues 55-58 were identified between the donor and acceptor heavy atoms of residues 55-58 listed above and Cys “SG” atoms as either donor or acceptor. The criteria for identifying H-bonds were distance between sulphur and heavy atoms below 4.1 Å and distance between donor hydrogen and sulphur or other heavy atoms below 3.2 Å ^80^.

### Data availability

Cx31.3 and Cx26 structures used in our templates are freely available in the worldwide PDB (code 6l3t and code 2zw3; *cf.* SI Table S1). Raw molecular dynamics data supporting the findings of the manuscript can be downloaded at http://downloadables.ttk.hu/heja/ConnexinMD2023.

### Code availability

Matlab scripts for trajectory analysis were made publicly available at https://github.com/hejalaszlo/ConnexinMD.

## Acknowledgements

This work was supported by National Research, Development and Innovation Office grants VEKOP- 2.1.1-15-2016-00156 and OTKA K124558. MD runs were performed using the supercomputer facility of the Governmental lnformation Technology Development Agency (KIFU, http://kifu.gov.hu).

## Author contributions

L.H. was responsible for study guidance, design, model building, analyses of MD simulations as well as original scripts. Á.S. performed homology modelling and conducted MD simulations. J.K. provided the first draft and overall assistance to the design and execution of the work. All authors contributed to manuscript preparation.

## Additional information

Additional information and datasets supporting the findings of this study are available as a Supplementary Information (SI) file. The online version contains supplementary material available for this paper at https://doi.org/………….

**Correspondence** should be addressed to László Héja.

