## Supplementary Information for "Simulation of gap junction formation reveals critical role of cysteines in connexon coupling"

#### Supplementary Information (SI)

László Héja*, Ágnes Simon & Julianna Kardos

Institute of Organic Chemistry, Research Centre for Natural Sciences, Magyar tudósok körútja 2, 1117, Budapest, Hungary.

### Tables

**SI Table S1.** Structures of connexin proteins in the PDB database

| Entry ID | Structure Title | construct | Experimental method | Primary citation | Resolution (Ǎ) |
|---|---|---|---|---|---|
| 2ZW3 | Connexin-26 gap junction channel at 3.5 angstrom resolution | GJ channel | X-ray diffraction | Maeda, S. et al. Structure of the connexin 26 gap junction channel at 3.5 A resolution, Nature, 2009, **58**, 597-602. | 3.5 |
| 5ER7 | Connexin-26 bound to calcium | GJ channel | X-ray diffraction | Bennett, B.C. et al. An electrostatic mechanism for Ca(2+)-mediated regulation of gap junction channels. Nat. Commun., 2016, **7**, 8770-8770 | 3.3 |
| 5ERA | Human Connexin-26 (Calcium-free) | GJ channel | X-ray diffraction | Bennett, B.C. et al. An electrostatic mechanism for Ca(2+)-mediated regulation of gap junction channels. Nat. Commun., 2016, **7**, 8770-8770 | 3.8 |
| 6UVR | Human Connexin-26 (Neutral pH open conformation) | GJ channel | Electron microscopy | Khan, A.K. et al. A Steric "Ball-and-Chain" Mechanism for pH-Mediated Regulation of Gap Junction Channels. Cell Rep., 2020, **31**, 1-11 | 4.0 |
| 6UVS | Human Connexin-26 (Low pH open conformation) | GJ channel | Electron microscopy | Khan, A.K. et al. Steric "Ball-and-Chain" Mechanism for pH-Mediated Regulation of Gap Junction Channels. Cell Rep., 2020, **31**, 1-11 | 4.2 |
| 6UVT | Human Connexin-26 (Low pH closed conformation) | GJ channel | Electron microscopy | Khan, A.K. et al. Steric "Ball-and-Chain" Mechanism for pH-Mediated Regulation of Gap Junction Channels. Cell Rep., 2020, **31**, 1-11 | 7.5 |
| 6L3T | Human Cx31.3/GJC3 connexin hemichannel in the absence of calcium | HC | Electron microscopy | Lee, H.J. et al. Cryo-EM structure of human Cx31.3/GJC3 connexin hemichannel. Sci. Adv. 2020, **6**, eaba4996 | 2.3 |
| 6L3U | Human Cx31.3/GJC3 connexin hemichannel in the presence of calcium | HC | Electron microscopy | Lee, H.J. et al. Cryo-EM structure of human Cx31.3/GJC3 connexin hemichannel. Sci. Adv. 2020, **6**, eaba4996 | 2.5 |
| 6L3V | R15G mutant of human Cx31.3/GJC3 connexin hemichannel | HC | Electron microscopy | Lee, H.J. et al. Cryo-EM structure of human Cx31.3/GJC3 connexin hemichannel. Sci. Adv. 2020, **6**, eaba4996 | 2.6 |
| 7JJP | Sheep Connexin-50 at 1.9 angstroms resolution by CryoEM | GJ channel | Electron microscopy | Flores, J.A. et al. Connexin-46/50 in a dynamic lipid environment resolved by CryoEM at 1.9 angstrom. Nat. Commun., 2020, **11**, 4331-4331 | 1.9 |
| 7JKC | Sheep Connexin-46 at 1.9 angstroms | GJ | Electron | Flores, J.A. et al. Connexin-46/50 in a dynamic lipid | 1.9 |

| | | | | | |
|---|---|---|---|---|---|
| | resolution by CryoEM | channel | microscopy | environment resolved by CryoEM at 1.9 angstrom. Nat. Commun., 2020, **11**, 4331-4331 | |
| 7JLW | Sheep Connexin-50 at 2.5 angstroms resolution, Lipid Class 1 | GJ channel | Electron microscopy | Flores, J.A. et al. Connexin-46/50 in a dynamic lipid environment resolved by CryoEM at 1.9 angstrom. Nat. Commun., 2020, **11**, 4331-4331 | 2.5 |
| 7JM9 | Sheep Connexin-50 at 2.5 angstroms reoslution, Lipid Class 2 | GJ channel | Electron microscopy | Flores, J.A. et al. Connexin-46/50 in a dynamic lipid environment resolved by CryoEM at 1.9 angstrom. Nat. Commun., 2020, **11** ,4331-4331 | 2.5 |
| 7JMC | Sheep Connexin-50 at 2.5 angstroms resolution, Lipid Class 3 | GJ channel | Electron microscopy | Flores, J.A. et al. Connexin-46/50 in a dynamic lipid environment resolved by CryoEM at 1.9 angstrom. Nat. Commun., 2020, **11**, 4331-4331 | 2.5 |
| 7JMD | Sheep Connexin-46 at 2.5 angstroms resolution, Lipid Class 1 | GJ channel | Electron microscopy | Flores, J.A. et al. Connexin-46/50 in a dynamic lipid environment resolved by CryoEM at 1.9 angstrom. Nat. Commun., 2020, **11**, 4331-4331 | 2.5 |
| 7JN0 | Sheep Connexin-46 at 2.5 angstroms resolution, Lipid Class 2 | GJ channel | Electron microscopy | Flores, J.A. et al. Connexin-46/50 in a dynamic lipid environment resolved by CryoEM at 1.9 angstrom. Nat. Commun., 2020, **11**, 4331-4331 | 2.5 |
| 7JN1 | Sheep Connexin-46 at 2.5 angstroms resolution, Lipid Class 3 | GJ channel | Electron microscopy | Flores, J.A. et al. Connexin-46/50 in a dynamic lipid environment resolved by CryoEM at 1.9 angstrom. Nat. Commun., 2020, **11**, 4331-4331 | 2.5 |
| 7QEQ | Human Connexin 26 dodecamer at 90mmHg $PCO_2$, pH7.4 | GJ channel | Electron microscopy | Brotherton, D.H. et al. Conformational changes and $CO_2$ -induced channel gating in connexin26. Structure, 2022, **30**, 697-706 | 1.9 |
| 7QER | Human Connexin 26 dodecamer at 55mm Hg PCO2, pH7.4 | GJ channel | Electron microscopy | Brotherton, D.H. et al. Conformational changes and $CO_2$ -induced channel gating in connexin26. Structure, 2022, **30**, 697-706 | 2.9 |
| 7QET | Human Connexin 26 dodecamer at 20mmHg $PCO_2$, pH7.4 | GJ channel | Electron microscopy | Brotherton, D.H. et al. Conformational changes and $CO_2$ -induced channel gating in connexin26. Structure, 2022, 30: 697-706 | 2.1 |
| 7QEW | Human Connexin 26 class 2 hexamer at 90mmHg $PCO_2$, pH7.4 | GJ channel | Electron microscopy | Brotherton, D.H. et al. Conformational changes and $CO_2$ -induced channel gating in connexin26. Structure, 2022, **30**, 697-706 | 2.1 |
| 7QEY | Human Connexin 26 class 1 hexamer at 90mmHg $PCO_2$, pH7.4 | GJ channel | Electron microscopy | Brotherton, D.H. et al. Conformational changes and $CO_2$ -induced channel gating in connexin26. Structure, 2022, **30**, 697-706 | 2.0 |

| | | | | | |
|---|---|---|---|---|---|
| 7QES | Human Connexin 26 at 55mm Hg $PCO_2$, pH7.4: two masked subunits, class A | GJ channel | Electron microscopy | Brotherton, D.H. et al. Conformational changes and $CO_2$ -induced channel gating in connexin26. Structure, 2022, **30**, 697-706 | 2.6 |
| 7QEU | Human Connexin 26 at 55mmHg $PCO_2$, pH7.4: two masked subunits, class B | GJ channel | Electron microscopy | Brotherton, D.H. et al. Conformational changes and $CO_2$ -induced channel gating in connexin26. Structure, 2022, **30**, 697-706 | 2.7 |
| 7QEV | Human Connexin 26 at 55mm Hg $PCO_2$, pH7.4:two masked subunits, class D | GJ channel | Electron microscopy | Brotherton, D.H. et al. Conformational changes and $CO_2$ -induced channel gating in connexin26. Structure, 2022, **30**, 697-706 | 2.9 |
| 7QEO | Human Connexin 26 at 55mm Hg $PCO_2$, pH7.4: two masked subunits, class C | GJ channel | Electron microscopy | Brotherton, D.H. et al. Conformational changes and $CO_2$ -induced channel gating in connexin26. Structure, 2022, **30**, 697-706 | 2.9 |
| 7F92 | Connexin43/Cx43/GJA1 gap junction intercellular channel in LMNG/CHS detergents at pH ~8.0 | GJ channel | Electron microscopy | Lee, H.J. et al. Conformational changes of human Cx43/GJA1 gap junction channel visualized by cryo-EM. Nat. Commun., 2023, **14**, 931 | 3.1 |
| 7F93 | Connexin43/Cx43/GJA1 gap junction intercellular channel in nanodiscs with soybean lipids at pH ~8.0 | GJ channel | Electron microscopy | Lee, H.J. et al. Conformational changes in the human Cx43/GJA1 gap junction channel visualized using cryo-EM. Nat. Commun., 2023, **14**, 931 | 3.6 |
| 7F94 | C-terminal truncated connexin43/Cx43/GJA1 gap junction intercellular channel with two conformationally different hemichannels | GJ channel | Electron microscopy | Lee, H.J. et al. Conformational changes in the human Cx43/GJA1 gap junction channel visualized using cryo-EM. Nat. Commun., 2023, **14**, 931 | 3.6 |
| 7XQF | C-terminal truncated connexin43/Cx43/GJA1 gap junction intercellular channel in POPE/CHS nanodiscs | GJ channel | Electron microscopy | Lee, H.J. et al. Conformational changes in the human Cx43/GJA1 gap junction channel visualized using cryo-EM. Nat. Commun., 2023, **14**, 931 | 2.2 |
| 7XQD | Structure of C-terminal truncated connexin43/Cx43/GJA1 gap junction intercellular channel in POPE/CHS nanodiscs (C1 symmetry) | GJ channel | Electron microscopy | Lee, H.J. et al. Conformational changes in the human Cx43/GJA1 gap junction channel visualized using cryo-EM. Nat. Commun., 2023, **14**:931 | 2.7 |
| 7XQB | Connexin43/Cx43/GJA1 gap junction intercellular channel in POPE/CHS nanodiscs at pH ~8.0 | GJ channel | Electron microscopy | Lee, H.J. et al. Conformational changes in the human Cx43/GJA1 gap junction channel visualized using cryo-EM. Nat. Commun., 2023, **14**:931 | 3.0 |
| 7XQJ | Hemichannel-focused structure of C-terminal truncated connexin43/Cx43/GJA1 | HC (half-GJ | Electron microscopy | Lee, H.J. et al. Conformational changes in the human Cx43/GJA1 gap junction channel visualized using cryo-EM. | 4.0 |

| | | | | | |
|---|---|---|---|---|---|
| | gap junction intercellular channel in POPE nanodiscs (PLN conformation) | channel) | | Nat. Commun., 2023, **14**:931 | |
| 7XQI | Hemichannel-focused structure of C-terminal truncated connexin43/Cx43/GJA1 gap junction intercellular channel in POPE nanodiscs (FIN conformation) | HC (half-GJ channel) | Electron microscopy | Lee, H.J. et al. Conformational changes in the human Cx43/GJA1 gap junction channel visualized using cryo-EM. Nat Commun., 2023, **14**:931 | 3.7 |
| 7XQH | Hemichannel-focused structure of C-terminal truncated connexin43/Cx43/GJA1 gap junction intercellular channel in POPE nanodiscs (GCN-TM1i conformation) | HC (half-GJ channel) | Electron microscopy | Lee, H.J. et al. Conformational changes in the human Cx43/GJA1 gap junction channel visualized using cryo-EM. Nat Commun., 2023, **14**:931 | 3.8 |
| 7XQG | Hemichannel-focused structure of C-terminal truncated connexin43/Cx43/GJA1 gap junction intercellular channel in POPE nanodiscs (GCN conformation) | HC (half-GJ channel) | Electron microscopy | Lee, H.J. et al. Conformational changes in the human Cx43/GJA1 gap junction channel visualized using cryo-EM. Nat. Commun., 2023, **14**, 931 | 3.8 |
| 7XQ9 | Connexin43/Cx43/GJA1 gap junction intercellular channel in GDN detergents at pH ~8.0 | GJ channel | Electron microscopy | Lee, H.J. et al. Conformational changes in the human Cx43/GJA1 gap junction channel visualized using cryo-EM. Nat. Commun., 2023, **14**, 931 | 3.3 |
| 7Z1T | Connexin43 gap junction channel structure in digitonin | GJ channel | Electron microscopy | Qi, C. et al. Structure of the connexin-43 gap junction channel in a putative closed state (to be published) | 2.3 |
| 7Z22 | Connexin43 gap junction channel structure in nanodisc | GJ channel | Electron microscopy | Qi, C. et al. Structure of the connexin-43 gap junction channel in a putative closed state (to be published) | 3.0 |
| 7Z23 | Connexin43 hemichannel in nanodisc | HC | Electron microscopy | Qi, C. et al. Structure of the connexin-43 gap junction channel in a putative closed state (to be published) | 4.0 |
| 7XKI | Human Cx36/GJD2 (N-terminal deletion BRIL-fused mutant) gap junction channel in soybean lipids (D6 symmetry) | GJ channel | Electron microscopy | Lee, S.N. et al. Cryo-EM structures of human Cx36/GJD2 neuronal gap junction channel. Nat- Commun., 2023, 14, 1347. | 3.4 |
| 7XKT | Human Cx36/GJD2 (BRIL-fused mutant) gap junction channel in detergents at 2.2 Angstroms resolution | GJ channel | Electron microscopy | Lee, S.N. et al. Cryo-EM structures of human Cx36/GJD2 neuronal gap junction channel. Nat. Commun., 2023, **14**, 1347. | 2.2 |
| 7XNH | Human Cx36/GJD2 gap junction channel with pore-lining N-terminal helices in soybean lipids | GJ channel | Electron microscopy | Lee, S.N. et al. Cryo-EM structures of human Cx36/GJD2 neuronal gap junction channel. Nat. Commun., 2023, **14**, 1347. | 3.1 |
| 7XNV | Structurally hetero-junctional human Cx36/GJD2 gap junction channel in soybean lipids (C6 symmetry) | GJ channel | Electron microscopy | Lee, S.N. et al. Cryo-EM structures of human Cx36/GJD2 neuronal gap junction channel. Nat. Commun., 2023, **14**, 1347. | 3.4 |

| | | | | | |
|---|---|---|---|---|---|
| 8HKP | Structurally hetero-junctional human Cx36/GJD2 gap junction channel in detergents (C6 symmetry) | GJ channel | Electron microscopy | Lee, S.N. et al. Cryo-EM structures of human Cx36/GJD2 neuronal gap junction channel. Nat. Commun., 2023, **14**, 1347. | 3.6 |
| 7XKK | Human Cx36/GJD2 gap junction channel in detergents | GJ channel | Electron microscopy | Lee, S.N. et al. Cryo-EM structures of human Cx36/GJD2 neuronal gap junction channel. Nat. Commun., 2023, **14**, 1347. | 3.2 |
| 7ZXM | Cryo-EM structure of Connexin 32 gap junction channel | GJ channel | Electron microscopy | Qi, C. et al. Structures of connexin 32 channels suggest a link between hemichannel gating and disease-associated mutations (to be published) | 2.1 |
| 7ZXN | Cryo-EM structure of Connexin 32 gap junction channel | GJ channel | Electron microscopy | Qi, C. et al. Structures of connexin 32 channels suggest a link between hemichannel gating and disease-associated mutations (to be published) | 3.1 |
| 7ZXO | Cryo-EM structure of Connexin 32 gap junction channel | GJ channel | Electron microscopy | Qi, C. et al. Structures of connexin 32 channels suggest a link between hemichannel gating and disease-associated mutations (to be published) | 2.5 |
| 7ZXP | Cryo-EM structure of Connexin 32 R22G mutation gap junction channel | GJ channel | Electron microscopy | Qi, C. et al. Structures of connexin 32 channels suggest a link between hemichannel gating and disease-associated mutations (to be published) | 2.4 |
| 7ZXQ | Cryo-EM structure of Connexin 32 R22G mutation hemi channel | HC | Electron microscopy | Qi, C. et al. Structures of connexin 32 channels suggest a link between hemichannel gating and disease-associated mutations (to be published) | 3.5 |
| 7ZXT | Cryo-EM structure of Connexin 32 W3S mutation hemi channel | HC | Electron microscopy | Qi, C. et al. Structures of connexin 32 channels suggest a link between hemichannel gating and disease-associated mutations (to be published) | 2.9 |

**SI Table S2. Intra-subunit SC distances between EL1 and EL2 in the Cx43HC-HC model.** SC distances are calculated between atoms having the smallest distance during 100 ns MD run. Average, minimum and maximum distances correspond to average, minimum and maximum values measured on all 12 subunits during the whole MD run. Average data are presented as distance ± SD [Å].

| SC atom pairs | Average distance | Minimum distance | Maximum distance |
|---|---|---|---|
| 51A “O” - 201S “OG” | 4.59 ± 0.74 | 2.52 | 7.64 |
| 52F “CE2” - 200L “CD2” | 5.69 ± 1.14 | 3.04 | 10.23 |
| 52F “CD2” – 199F “O” | 3.71 ± 0.57 | 2.66 | 6.67 |
| 52F “CZ” – 198C(3) “CB” | 3.70 ± 0.31 | 2.96 | 9.14 |
| 53R “O” - 199F “N” | 2.81 ± 0.12 | 2.47 | 3.73 |
| 53R “O” – 198C(3) “CA” | 3.36 ± 0.17 | 2.81 | 4.82 |
| 54C(1) “SG” – 198C(3) “SG” | 2.03 ± 0.04 | 1.86 | 2.21 |
| 55N “OD1” – 199F “CE2” | 5.56 ± 1.14 | 2.76 | 13.18 |
| 55N “OD1” – 198C “N” | 7.59 ± 0.97 | 3.13 | 10.11 |
| 55N “N” – 197D “O” | 3.09 ± 0.49 | 2.51 | 6.13 |
| 55N “O” -196V “CG2” | 5.28 ± 1.47 | 2.75 | 10.85 |
| 58Q “NE2” – 193P “O” | 8.39 ± 1.11 | 2.75 | 13.33 |
| 58Q “OE1” – 192C “SG” | 7.00 ± 1.38 | 2.82 | 11.80 |
| 56T “OG1” – 196V “CG2” | 5.01 ± 0.89 | 2.86 | 8.38 |
| 69S “OG” - 187C(1) “CB” | 7.56 ± 1.05 | 3.71 | 10.33 |
| 68K “CD” – 187C(1) “SG” | 6.31 ± 0.99 | 3.23 | 9.67 |
| 65C(3) “SG” – 189R “CB” | 5.62 ± 0.78 | 3.34 | 8.62 |
| 65C(3) “SG” – 192C(2) “SG” | 4.11 ± 1.10 | 2.99 | 7.92 |
| 64V “CG2”- 191P “CD” | 4.21 ± 0.53 | 3.18 | 7.69 |
| 64V “CG2” – 192 C(2) “SG” | 5.27 ± 1.21 | 3.08 | 10.05 |
| 61C(2) “SG” – 192 C(2) “SG” | 2.03 ± 0.04 | 1.85 | 2.19 |

### Figures

**A**

```
Cx43   1   MGDWSALGKLLDKVQAYSTAGGKVWLSVLFIFRILLLGTAVESAWGDEQSAFRCNTQQPGCENVCYDKSFPISHVRFWVL  80
Cx31   1   ------LRRLLAEESRRSTPVGRLLLPVLLGFRLVLLAASGPGVYGDEQSEFVCHTQQPGCKAACFDAFHPLSPLRFWVF  79

Cx43  81   QIIFVSVPTLLYLAHVFY----------------------(52)---------------------------LLRTYIISI 159
Cx31  80   QVILVAVPSALYMGFTLY----------------------(35)---------------------------LLWAYVAQL 141

Cx43 160   LFKSIFEVAFLLIQWYIYGFSLSAVYTCKRDPCPHQVDCFLSRPTEKTIFIIFMLVVSLVSLALNIIELFYVFFKGVKDR 239
Cx31 142   GARLVLEGAALGLQYHLYGFQMPSSFACRREPCLGSITCNLSRPSEKTIFLKTMFGVSGFCLLFTFLELV---------- 211

Cx43 240   VKGKSDPYHATSGALSPAKDCGSQKYAYFNGCSSPTAPLSPMSPPGYKLVTGDRNNSSCRNYNKQASEQNWANYSAEQNR 319
Cx31       --------------------------------------------------------------------------------
```

**B**

```
Cx43   1   MGDWSALGKLLDKVQAYSTAGGKVWLSVLFIFRILLLGTAVESAWGDEQSAFRCNTQQPGCENVCYDKSFPISHVRFWVL  80
Cx26   1   -MDWGTLQTILGGVNKHSTSIGKIWLTVLFIFRIMILVVAAKEVWGDEQADFVCNTLQPGCKNVCYDHYFPISHIRLWAL  79

Cx43  81   QIIFVSVPTLLYLAHVFY----------------------(52)---------------------------LLRTYIISI 159
Cx26  80   QLIFVSTPALLVAMHVAY----------------------(33)---------------------------LWWTYTSSI 140

Cx43 160   LFKSIFEVAFLLIQWYIY-GFSLSAVYTCKRDPCPHQVDCFLSRPTEKTIFIIFMLVVSLVSLALNIIELFYVFFKGVKD 238
Cx26 141   FFRVIFEAAFMYVFYVMYDGFSMQRLVKCNAWPCPNTVDCFVSRPTEKTVFTVFMIAVSGICILLNVTELCYLLIRYCSG 220

Cx43 239   RVKGKSDPYHATSGALSPAKDCGSQKYAYFNGCSSPTAPLSPMSPPGYKLVTGDRNNSSCRNYNKQASEQNWANYSAEQN 318
Cx26 221   KSKKP----------------------------------------------------------------------------

Cx43 319   RMGQAGSTISNSHAQPFDFPDDNQNSKKLAAGHELQPLAIVDQRPSSRASSRASSRPRPDDLEI
Cx26       ---------------------------------------------------------------
```

**SI Figure S1. (A)** Sequence alignment between Cx43 and Cx31.3 (PDB code: 6l3t) [1] generated by Swiss-Model. Transmembrane (TM) amino acids (AA) of Cx43 and Cx31.3 TM regions according to PPM are shown in red. TM AAs of Cx43 are as follows: 20-46, 74-94, 154-176, 205-226. **(B)** Sequence alignment between Cx43 and Cx26 (PDB code: 2zw3) [2] generated by Swiss-Model. TM amino acids (AA) of Cx43 and Cx26 TM regions according to PPM are shown in red. TM AAs of Cx43 are as follows: 20-46, 74-94, 154-176, 204-230.

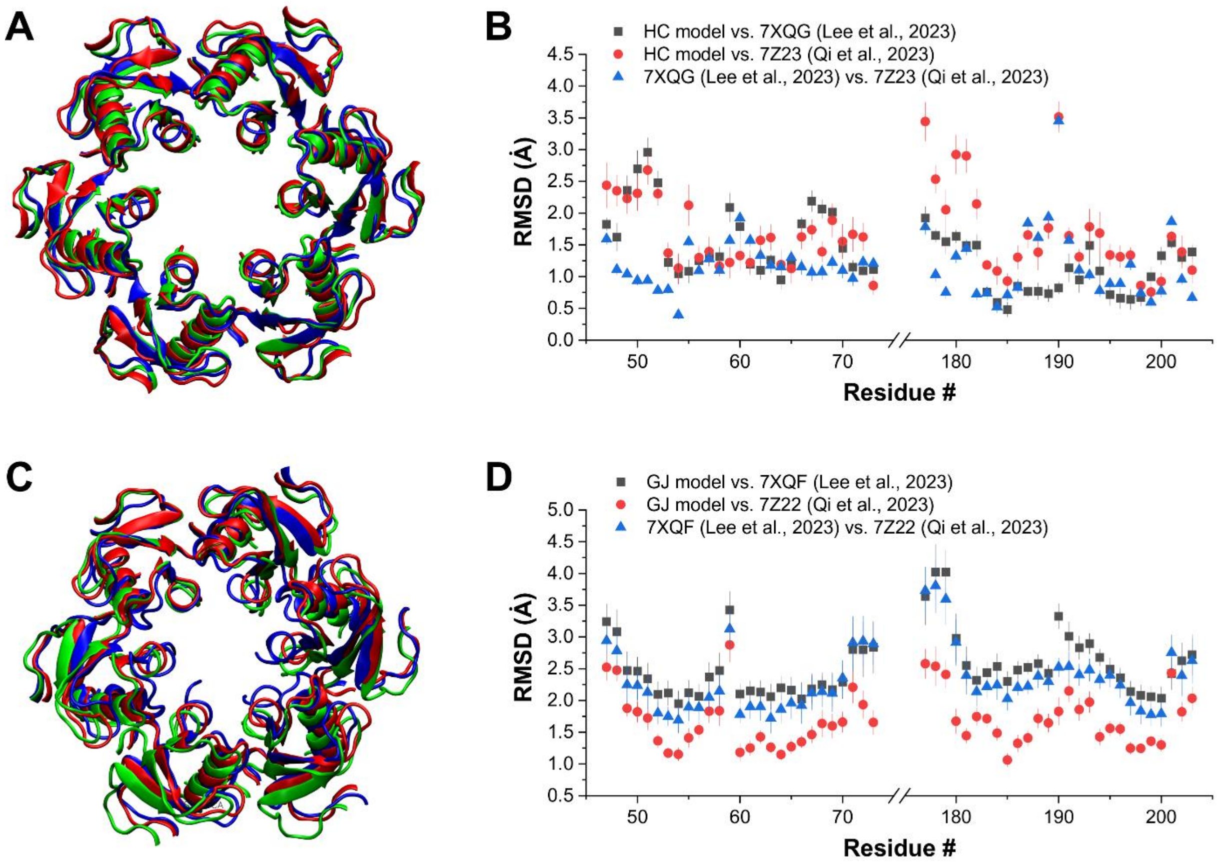


**SI Figure S2. Comparison of Cx43 models to experimental Cx43 structures. (A)** 3D alignment of the extracellular regions of the Cx43HC model in the closed disulphide configuration (closed Cx43HC model, this work) at the beginning of the 100 ns MD run (blue) with the experimental Cx43 HC structure 7Z23 [3] (red) and the experimental half-GJ structure 7XQG [4] (green). **(B)** RMSD difference of individual extracellular residues between the closed Cx43HC model (this work), the experimental Cx43 HC structure 7Z23 [3] and the experimental half-GJ structure 7XQG [4]. Data are presented as mean ± SEM, averaged over all subunits. **(C)** 3D alignment of the extracellular regions of the Cx43GJ model in the closed disulphide configuration (closed Cx43GJ model, this work) at the beginning of the 100 ns MD run (blue) with the experimental Cx43 GJ structures 7Z22 [3] (red) and 7XQF [4] (green). For clarity, only the A-F subunits are shown. **(D)** RMSD difference of individual extracellular residues between the closed Cx43HC model (this work) and the experimental Cx43 GJ structures 7Z22 [3]and 7XQF [4]. Data are presented as mean ± SEM, averaged over all subunits.

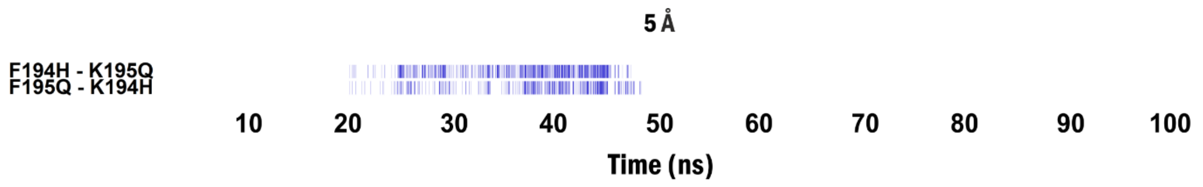


**SI Figure S3. Appearance of trans-GJ SCs in the Cx43HC-HC model after moving HCs away to 5 Å distance.** Presence of specific trans-GJ SCs during 100 ns MD simulation show only a short appearance of trans-GJ interactions after moving HCs away to 5 Å distance. Trans-GJ SC pairs are designated as subunit name + residue number + 1-letter residue name.

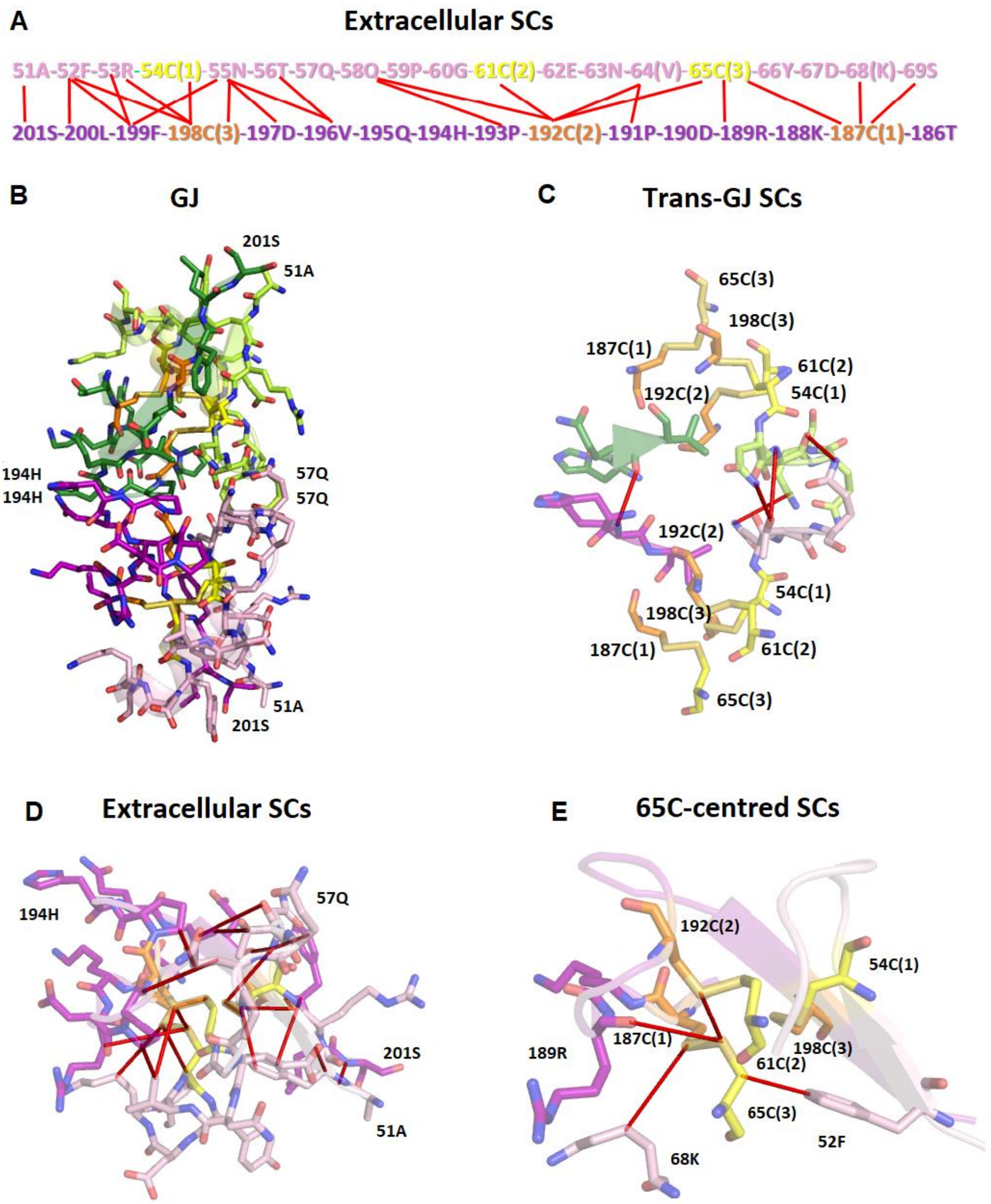


**SI Figure S4. Stabilization centers (SCs) in the Cx43HC-HC model are located near to extracellular cystines.** (**A**) Two-dimensional scheme of SC patterns shaped by inter-loop cystine disulphide bonds. Red lines indicate SC residue contacts as characterized by minimum atomic distance in Å (brackets) (*see* Methods): 51A-201S(2.52); 52F-200L(3.04); 52F‒199F(2.66); 52F‒198C(3)(2.96); 53R‒199F(2.47); 53R-198C(3)(2.81); 54(1)‒198(3)(1.86); 55N-199F(2.76); 55N-198C(3)(3.13); 55N-197D(2.51); 55N-196V(2.75); 56T-196V(2.45); 58Q‒193P(2.75); 58Q‒192C(2)(2.82);

69S-187C(1)(3.71); 68K-187C(1)(3.23); 65C(3)-189R(3.34); 65C(3)-192C(2)(2.99); 64V-191P(3.18); 64V-192C(2)(3.08); 61C(2)-192C(2(1.85) (*see* Methods). (**B**) The quasi-mirrored three-dimensional configuration of the Cx43HC-HC model. (**C**) The mid trans-GJ pattern of Cx43HC-HC model comes into view as coupled pairs of pre/post-GJ EL1 55N-56T-57Q along with EL2 194H-195Q-196V (trans-GJ SCs) plus trans-GJ H-bonding formation between pre GJ EL1 58Q and post EL1 55N (or *vice versa*) associated with cystine disulphide bonds located in heads of pyramids-like silhouettes. Trans-GJ SCs and H-bonding interactions are indicated by red lines. (**D**) The three-dimensional pattern of extracellular cystine disulphide centred GJ SC pattern is characterized by red lines. (**E**) The unique EL1 65C(3)-centred three-dimensional SC pattern combines EL1 52F - 65C(3) in addition to EL1-EL2 65C(3) - 189R, 65C(3) - 192C(2) and 68K - 187C(1). Intra-loop (EL1) and inter-loop (EL1-EL2) interactions are shown by red lines. Colour codes: yellow - EL1 54C(1), 61C(2), 65C(3); orange – EL2 187C(1), 192(2), 198C(3); lemon – pre-GJ EL1 residues; Forest – pre-GJ EL2 EL1 residues; pink – post-GJ EL1 residues; purple – post-GJ EL2 residues.

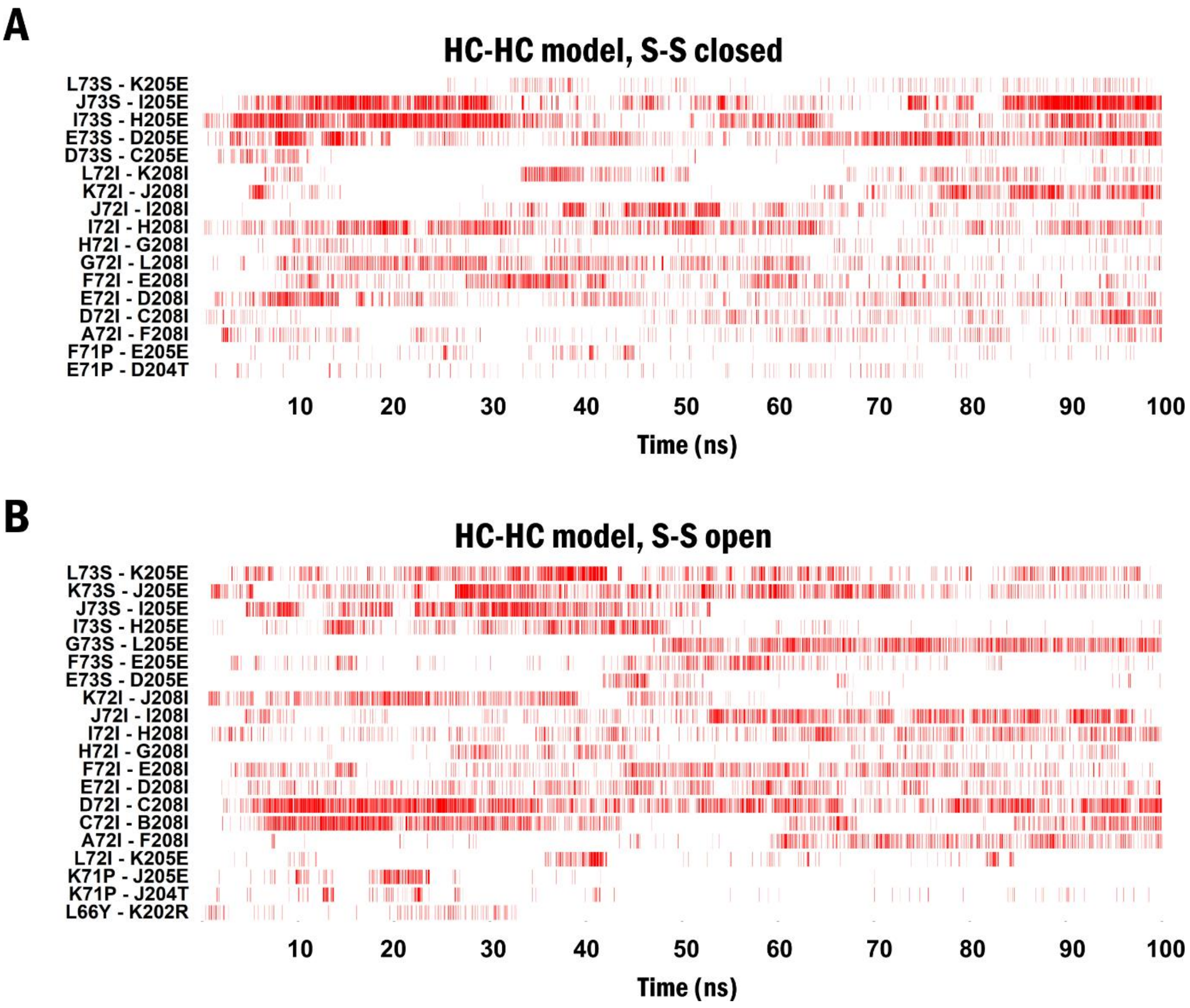


**SI Figure S5. Inter-subunit stabilization centers (SCs) within HCs are not disrupted by opening of disulphide bonds in the Cx43 HC-HC model.** (**A**) Presence of specific inter-subunit SCs during 100 ns MD simulation in the Cx43 HC-HC model in the closed disulphide configurations. (**B**) Presence of specific inter-subunit SCs during 100 ns MD simulation in the Cx43 HC-HC model in the open disulphide configurations. Inter-subunit SC pairs are designated as subunit name + residue number + 1-letter residue code. Only SCs formed between two connexin subunits in the same HC are shown.
